# Stability of extemporaneously compounded amiloride nasal spray

**DOI:** 10.1101/2020.04.16.044586

**Authors:** Venkata Yellepeddi, Casey Sayre, Anna Burrows, Kevin Watt, Simon Davies, John Strauss, Marco Battaglia

**Author notes:** Corresponding author (VY).

## Abstract

Anxiety disorders (AD) are the most common mental illnesses affecting an estimated 40 million adults in the United States. Amiloride, a diuretic agent, has shown efficacy in treating AD in preclinical models by inhibiting the acid-sensing ion channels (ASIC). By delivering amiloride via nasal route, rapid onset of action can be achieved due to direct “nose-to-brain” access. Therefore, this study reports the formulation, physical, chemical, and microbiological stability of an extemporaneously prepared amiloride 2 mg/mL nasal spray. The amiloride nasal spray was prepared by adding 100 mg of amiloride hydrochloride to 50 mL of sterile water for injection in a sterile reagent bottle. A stability-indicating high-performance liquid chromatography (HPLC) method was developed and validated. Forced-degradation studies were performed to confirm the ability of the HPLC method to identify the degradation products from amiloride distinctively. The physical stability of the amiloride nasal spray was assessed by pH, clarity, and viscosity assessments. For chemical stability studies, samples of nasal sprays stored at room temperature were collected at time-points 0, 3 hr., 24 hr., and 7 days and were assayed in triplicate using the stability-indicating HPLC method. Microbiological stability of the nasal spray solution was evaluated for up to 7 days based on the sterility test outlined in United States Pharmacopoeia (USP) chapter 71. The stability-indicting HPLC method identified the degradation products of amiloride without interference from amiloride. All tested solutions retained over 90% of the initial amiloride concentration for the 7-day study period. There were no changes in color, pH, and viscosity in any sample. The nasal spray solutions were sterile for up to 7 days in all samples tested. An extemporaneously prepared nasal spray solution of amiloride hydrochloride (2 mg/mL) was physically, chemically, and microbiologically stable for 7 days when stored at room temperature.

## Introduction

Anxiety disorders (AD) are the most common mental illnesses in every age group, affecting 25% of children and an estimated 40 million adults in the United States [1]. Risk factors for developing AD include genetics, life adversities, and subtle brain chemistry alterations [1].

Pharmacological and cognitive-behavioral interventions, alone or in combination, are typically employed to treat AD [2, 3]. Contemporary first-line pharmacological agents for AD include selective serotonin reuptake inhibitors (SSRIs) and some serotonin noradrenaline-reuptake inhibitors (SNRIs). SSRIs and SNRIs are better tolerated than older generation tricyclic antidepressants and show moderate-to-good effectiveness. However, the effectiveness and duration of treatment are not significantly different compared with tricyclic antidepressants, and many people experience relapse [4]. This highlights the unmet need for improved therapeutics in AD [2, 3].

In a series of comparative human and preclinical studies of responses to CO_2_ (an unconditioned stimulus that evokes panic-like responses in humans at risk for panic disorder and hyperventilation in rodents) [5-7], we reported that life adversities enhance the liability to anxiety and pain through the enrichment of acid-sensing ion channel (ASIC) genes -1 and -2 [8, 9]. Coherent with these findings, ASIC-antagonist amiloride (nebulized to bypass the blood-brain barrier) normalizes the enhanced anxious and nociceptive responses that are proper of mice exposed to early disrupted maternal care and the nociceptive responses of rats that underwent prenatal maternal stress [9]. Taken together, these data suggest that amiloride has the potential to provide effective treatment in humans both in certain AD and for some conditions characterized by pain.

Amiloride hydrochloride is a pyrazine-carbonyl-guanidine [10, 11] salt of a moderately strong base (pKa 8.7), and an antikaliuretic-diuretic agent. Amiloride is currently approved by the Food and Drugs Administration (FDA) as adjunctive treatment with thiazide diuretics or other kaliuretic diuretic agents in congestive heart failure, or hypertension [10]. Amiloride is available only as a tablet formulation for oral administration with an onset of action time of 2 hours, with peak plasma levels reached within 3 to 4 hours [10, 11]. This time to onset of action is too slow for panic attacks, which have a very rapid onset. However, rapid onset of action can be achieved via the intranasal route of administration that allows for rapid absorption of drugs into the central nervous system (CNS) via the “nose-to-brain” route [12, 13]. Intranasal administration also provides a non-invasive point of access into the CNS and reduces the risk of needle-stick injuries due to parenteral administration in hospital and emergency department settings [14]. Furthermore, intranasal administration allows for simple, self-administration, which facilitates patients’ adherence [15].

Because amiloride is an FDA-approved drug, we can extemporaneously compound amiloride nasal spray for the treatment of AD. However, currently, there are no reports on the formulation and stability of amiloride nasal spray that can be adapted by the compounding pharmacists to prepare amiloride nasal spray. Therefore, the objective of this study is to develop amiloride nasal spray formulation and report its stability at room temperature. In this study, we report on the extemporaneous formulation of amiloride nasal spray and its physical, chemical, and microbiological stability.

## Materials and Methods

### Materials

Amiloride hydrochloride powder was purchased from EMD Millipore Corporation, Temecula, CA. Lot. 3224630. The United States Pharmacopeia (USP) reference standard of amiloride hydrochloride was purchased from, USP Convention, Rockville, MD. Lot. R052W0. The USP reference standard was used for the HPLC method development and validation. The USP grade sterile water for injection used for the preparation of amiloride nasal spray was purchased from RMBIO, Missoula, MT, USA. HPLC grade glacial acetic acid and methanol were purchased from EMD Millipore Corporation, Temecula, CA, USA. The chemicals and reagents used for the forced degradation studies were purchased from Millipore Sigma, St. Louis, MO, USA. The reversed-phase C18 HPLC column used was Waters Nova-Pak^®^ purchased from Waters Corporation, Milford, MA. Lot. 1130380642.

### Extemporaneous preparation of amiloride nasal spray

Amiloride nasal spray was formulated by dissolving 20 mg of amiloride hydrochloride in 10 mL sterile water for injection (2 mg/mL) in a laminar flow hood. The sterile water for injection was previously heated to 40°C using a water bath. Whenever possible, amiloride was protected from light to avoid photodegradation. The formulation was filtered using 0.22 µm nylon syringe filters into 10-mL sterile syringes in a laminar-airflow hood, Labconco, Logic^+^ A2, Kansas City, MO. Details of the steps involved in the procedure are provided as S1 Appendix in the supporting information.

### Stability-indicating high-performance liquid chromatography method

The stability-indicating high-performance liquid chromatography (HPLC) method used to analyze amiloride and its degradation products was developed and validated according to FDA guidelines for bioanalytical method validation [16]. Briefly, the mobile phase consisted of a 25:75 ratio of methanol: water with the final pH adjusted to 3.6 with glacial acetic acid. Amiloride was detected using a photodiode array detector at a maximum wavelength of 284 nm. The chromatographic separation was achieved using a 5-µm particle size, 3.9 mm × 15 cm L1 column. The runs for each injection involved a gradient elution programmed, as shown in Table 1.

**Table 1.** HPLC gradient elution flow program for analysis of amiloride.

| Time (min) | Flow rate (mL/min) | % Water | % Methanol |
| --- | --- | --- | --- |
| 0.00 | 1 | 75 | 25 |
| 21.00 | 1 | 75 | 25 |
| 22.09 | 1 | 100 | - |
| 26.10 | 1 | 100 | - |
| 27.99 | 1 | 75 | 25 |

HPLC analysis was performed by injecting 20 µL of the amiloride sample into the separation module equipped with a photodiode array detector, PDA detector, Model No.2998, Waters Corporation, Milford, MA. Data acquisition and analysis were performed using Empower; version 3 (Waters Corp., Milford, MA). The HPLC method was validated according to the International Council on Harmonisation (ICH) guidelines for linearity, accuracy and precision, robustness, and ruggedness [17]. A standard 5-point calibration curve was constructed by linear regression of the peak areas of the amiloride peak obtained from amiloride hydrochloride USP reference standard solutions at concentrations 12.5, 25, 50, 100, and 200 µg/mL (*r*^*2*^ = 0.9998).

The specificity of the HPLC method to degradation products of amiloride was assessed by subjecting amiloride to forced-degradation conditions including hydrolysis, oxidation, photodegradation, thermal degradation, and ultraviolet (UV) irradiation. Stability indicating forced-degradation studies involved treatment of 100 µg/mL amiloride hydrochloride solution with 0.1M hydrochloric acid for 1 hour to assess for acid hydrolysis, 0.1M sodium hydroxide for 1 hour to evaluate for alkaline hydrolysis, 0.1M hydrogen peroxide overnight to determine for oxidation, UV radiation (UV light of laminar-airflow hood) overnight to assess for photostability, and temperature of 60°C (using a laboratory hot plate) overnight to determine for thermal stability studies.

### Chemical stability studies

For the chemical stability analysis, a batch of 20 amiloride nasal sprays (2 mg/mL) were prepared as described earlier. Five syringes were randomly selected from the batch and placed on a dry ventilated surface (mean ± S.D. temperature of 22.7 ± 0.8 °C and relative humidity [RH] of 32.5% ± 5%). At 0 (immediately after preparation), 3 hours, 24 hours, and 7 days, a pipette was used to transfer a 100-µL sample of nasal spray into 15 mL centrifuge tube. The solution was diluted to obtain a final concentration of 200 µg/mL. The final solution was injected into HPLC in triplicate for analysis. The percentage assay values for stability samples were calculated using the calibration curve described above, using peak areas of amiloride obtained after integrating peaks from chromatograms of stability samples.

### Physical stability tests

Color, visual clarity, pH, and viscosity were also evaluated at 0, 3 hours, 24 hours, and 7 days. The samples were visually inspected against black and white backgrounds using a high-intensity lamp at each time point to evaluate the characteristics of color and clarity. The pH meter, Seven Easy, Model No. S20, Mettler Toledo, Columbus, OH, was calibrated with standard buffer solutions of pH 4, 7, and 10, was used for pH analysis. Viscosity was measured using a rheometer, Kinexus ultra^+^ rheometer, Malvern PANalytical, Malvern, UK.

### Microbiological stability analysis

For microbiological stability analysis, 20 syringes of amiloride nasal spray, prepared as described above, were placed on a ventilated surface at room temperature (median ± S.D. temperature of 22.7 ± 0.8 °C and RH of 32.5% ± 5%). Microbiological stability was evaluated at 0 and 7 days after storage at room temperature. To ascertain microbiological stability, samples were subjected to sterility tests described in USP Chapter 71 [18]. The sterility test was carried out using the membrane filtration technique with appropriate negative controls. Briefly, the sample was hand filtered across two separate filters, followed by the addition of tryptic soy broth medium (TSB) to one filter and fluid thioglycollate medium (FTM) to the other. The filter with TSB medium was incubated at 20 - 25°C and the filter with FTM was incubated at 30-35 °C for over 2 weeks. During these 2 weeks, the media were examined for macroscopic evidence of microbiological growth. Data are presented as the presence or absence of microbial growth as determined by visual examination. All microbiological analyses were performed at Compounder’s International Analytical Laboratory, Castle Rock, Colorado, USA.

### Data analysis

The stability was defined as the retention of at least 90% of the initial concentration of amiloride nasal spray. The chemical stability experiments were performed in triplicate after 3 samples were randomly collected from 20 nasal sprays. Data were represented as the mean ± S.D. percent of the initial concentrations remaining. For pH, particulate matter, and viscosity tests, differences between samples at different time points were compared using 1-way analysis of variance (ANOVA). The *a priori* level of significance was 0.05. Statistical analyses were performed using GraphPad Prism, Version 8, GraphPad Software, San Diego, CA. For sterility testing, three samples from each time point (0 and 7 days) were filtered in triplicate for each medium.

## Results and discussion

The amiloride nasal spray solutions were successfully prepared and were colorless with no visible particulate matter when inspected against dark and light backgrounds. The validated HPLC method showed that amiloride was eluted at ∼ 10.3 minutes. A freshly prepared solution of amiloride hydrochloride was analyzed using the HPLC method and compared with the USP reference standard of amiloride hydrochloride. The retention time and the shape of the peaks were similar between the sample and the standard solutions. The mean ± S.D. percentage assay of amiloride hydrochloride was 96.8% ± 0.4%. The chromatograms showed sharp and distinct peaks for each analyte without interference or co-elution from the contents of the mobile phase (Fig 1).

**Fig 1.**
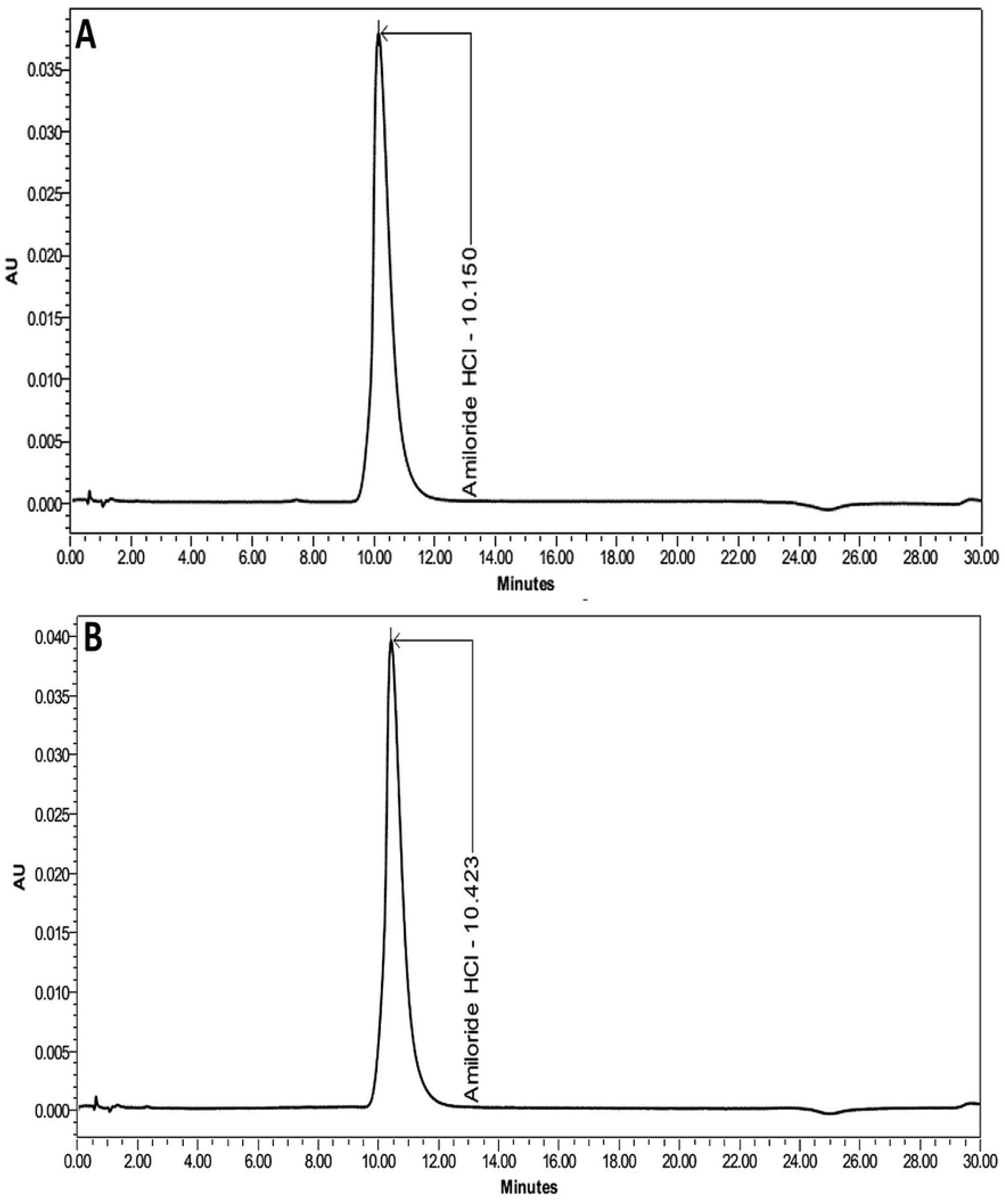
Amiloride hydrochloride pure drug and reference standard chromatograms. Chromatograms representing 0.1 mg/mL amiloride hydrochloride solutions in sterile water for injection. A) 0.1 mg/mL amiloride hydrochloride USP reference standard solution. B) 0.1 mg/mL amiloride hydrochloride solution used for the preparation of nasal spray.

Relative S.D. for replicate injections was 0.4% (USP limit of ≤2%), indicating the suitability of the HPLC method for the assay of amiloride nasal spray. The standard curve of amiloride nasal spray was linear over the range of concentrations (12.5 to 200 µg/mL, *r*^*2*^ = 0.999). The chromatograms representing amiloride at various concentrations used for the standard curve are provided as Fig 2. Results from the validation studies showed that the parameters accuracy, precision, robustness, and ruggedness were within the specified limits outlined in ICH guidelines (S2 appendix) [17].

**Fig 2.**
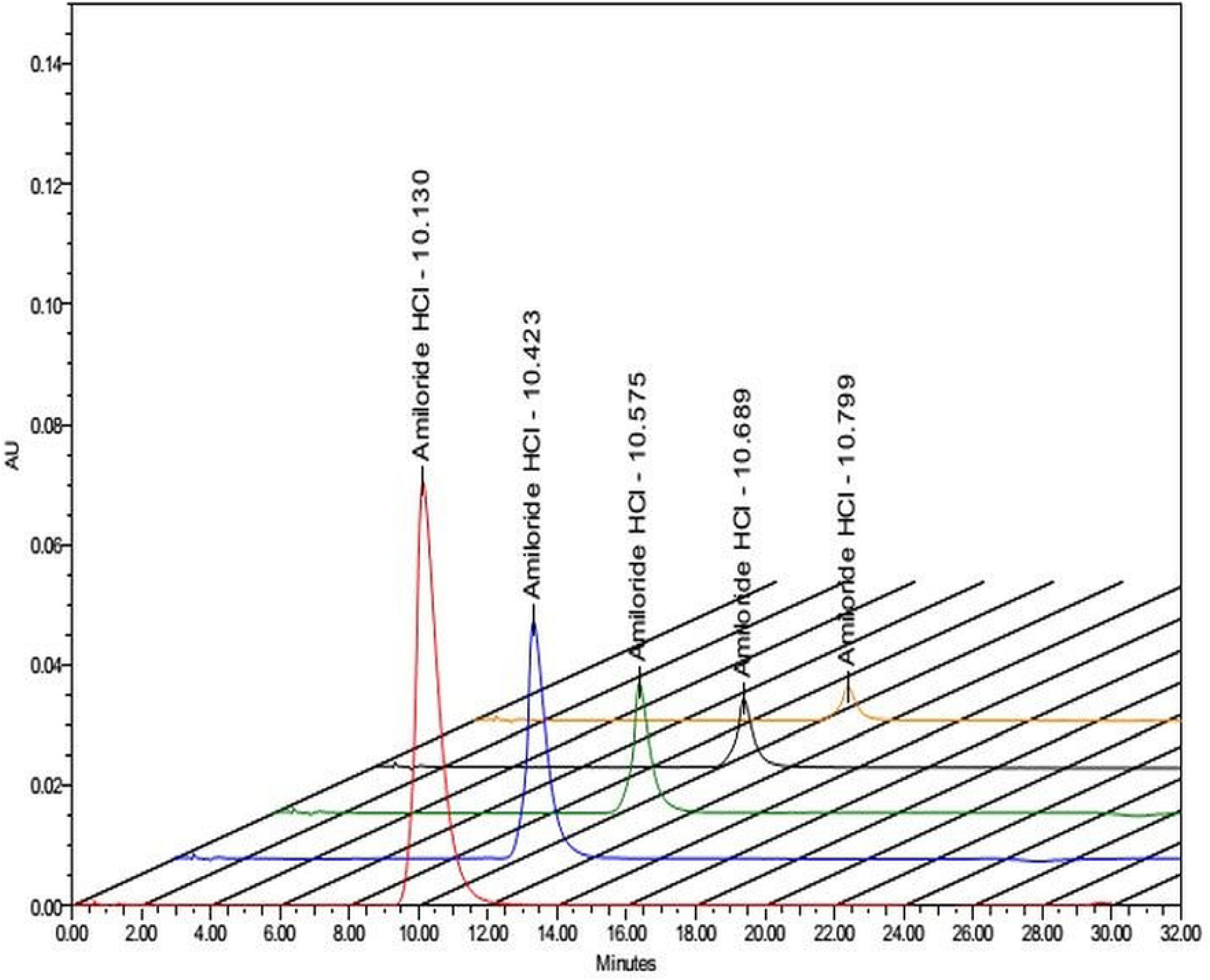
Overlay of chromatograms representing standard curve of amiloride. From left to right, chromatograms represent amiloride concentrations 200, 100, 50, 25, and 12.5 µg/mL.

Hydrolysis and photodegradation are the major degradation pathways of amiloride [19, 20]. Therefore, it is essential that the HPLC method can identify the degradation products of amiloride. The forced degradation studies were performed to assess the capability of the HPLC method to identify the degradation products from intact amiloride distinctly. The chromatograms of amiloride samples subjected to hydrolysis and photodegradation identified the degradation products of amiloride. These results indicate that the HPLC method used has the potential to identify major degradation products of amiloride, and can be successfully used for stability studies of amiloride nasal spray.

The amiloride 2-mg/mL nasal spray stored at room temperature showed good physical and chemical stability for up to 7 days (Table 2). The percentage of the initial amiloride concentration remaining at 7 days was 98.7%, indicating satisfactory chemical stability of amiloride nasal spray (Fig 3). However, due to its potential for photodegradation, caution must be exercised by compounding pharmacists to avoid exposing amiloride to light. Furthermore, the nasal spray must be dispensed in containers that preserve amiloride by protection from photodegradation. The data from the physical stability studies (pH and viscosity) showed no significant differences among the samples at time points 0, 3 hours, 24 hours, and 7 days (*p* > 0.05).

**Table 2.** Stability of 2 mg/mL amiloride nasal spray solution.

| **Stability Parameter Analyzed** | **0** | **3 hours** | **24 hours** | **7 days** |
| --- | --- | --- | --- | --- |
| **% Assay of amiloride<sup>a</sup>** | 96.51 ± 0.34 | 96.64 ± 0.19 | 95.24 ± 0.1 | 98.23 ± 0.44 |
| **pH<sup>b</sup>** | 4.47 ± 0.23 | 4.52 ± 0.16 | 4.51 ± 0.33 | 4.54 ± 0.26 |
| **Viscosity (cP)<sup>c</sup>** | 0.89 ± 0.04 | 0.91 ± 0.02 | 0.9 ± 0.01 | 0.94 ± 0.01 |

<sup>a</sup> Values represent mean ± S.D. after triplicate analysis of three samples obtained from three nasal sprays. The % assay values are calculated from a calibrated curve prepared from a standard solution of USP amiloride hydrochloride.
<sup>b</sup> Values represent mean ± S.D. pH after triplicate analysis from three nasal sprays. The pH reported is the final pH of the nasal spray after addition of amiloride hydrochloride solution to water. No significant difference in pH was observed between samples at various time points based on results from ANOVA ( $p > 0.05$ ).
<sup>c</sup> Values represent mean ± S.D. after triplicate analysis from three nasal sprays. No significant difference in viscosity was observed between samples at various time points based on results from ANOVA ( $p > 0.05$ ).

**Fig 3.**
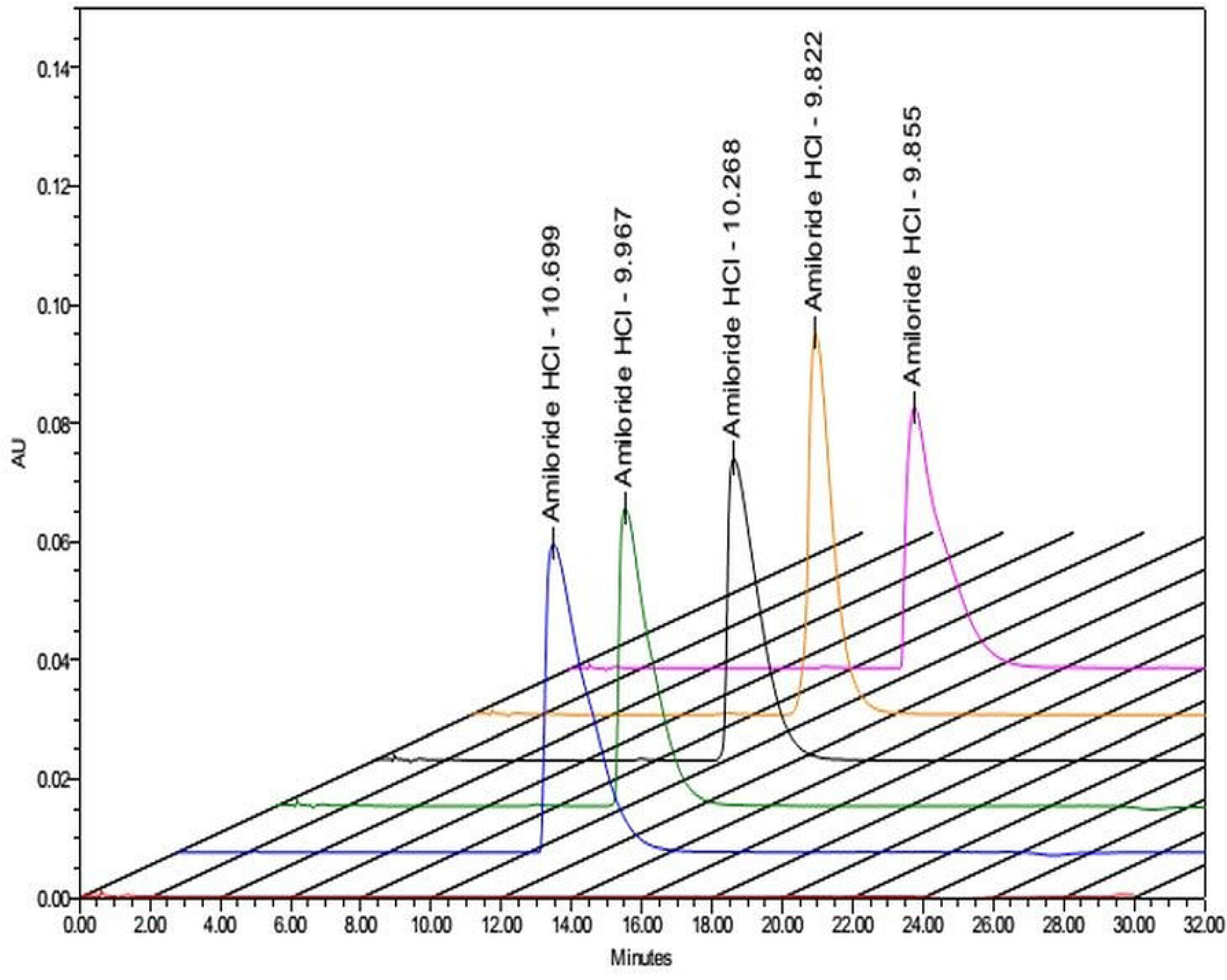
Chemical stability of amiloride nasal spray. Overlay of chromatograms representing amiloride nasal spray samples assayed after storage in bottles at room temperature. From left to right, chromatograms represent USP reference standard, 0, 3 hrs, 24 hrs, and 7 days samples of 0.2 mg/mL amiloride hydrochloride solutions.

FDA requires that all nasal spray solutions for human use must be free of microbiological contamination due to their direct interaction with mucosal surfaces [21]. Therefore, we evaluated the microbiological stability of amiloride nasal spray by confirming its sterility after storage at room temperature for 7 days using the method outlined in USP 71. The results from this testing revealed that the extemporaneously prepared amiloride nasal spray was microbiologically stable for up to 7 days. In all samples, tested, there were no signs of microbiological growth over two weeks.

## Conclusions

An extemporaneously prepared nasal spray solution of amiloride hydrochloride 2 mg/mL exhibited physical, chemical, and microbiological stability over 7 days when stored at room temperature in sterile syringes.

## Acknowledgements

We would like to thank Dr. Hamid Ghandehari for providing access to the rheometer in his laboratory.

## Supporting information

**S1 Appendix. Formulation of amiloride nasal spray**. Document with step-by-step directions for extemporaneously compounding amiloride nasal spray 2 mg/mL.

**S2 Appendix. HPLC method validation results**. Excel file containing HPLC method validation data for amiloride hydrochloride.

## References

1. Facts & Statistics. Anxiety and Depression Association of America (https://adaa.org/about-adaa/press-room/facts-statistics [cited 2020 01/09/2020].

2. Bystritsky A, Khalsa SS, Cameron ME, Schiffman J. Current diagnosis and treatment of anxiety disorders. P T. 2013;38(1):30-57. Epub 2013/04/20. PubMed PMID: 23599668; PubMed Central PMCID: PMCPMC3628173.

3. Farach FJ, Pruitt LD, Jun JJ, Jerud AB, Zoellner LA, Roy-Byrne PP. Pharmacological treatment of anxiety disorders: current treatments and future directions. J Anxiety Disord. 2012;26(8):833-43. Epub 2012/10/02. doi: 10.1016/j.janxdis.2012.07.009. PubMed PMID: 23023162; PubMed Central PMCID: PMCPMC3539724.

4. Bruce SE, Yonkers KA, Otto MW, Eisen JL, Weisberg RB, Pagano M, et al. Influence of psychiatric comorbidity on recovery and recurrence in generalized anxiety disorder, social phobia, and panic disorder: a 12-year prospective study. Am J Psychiatry. 2005;162(6):1179-87. Epub 2005/06/03. doi: 10.1176/appi.ajp.162.6.1179. PubMed PMID: 15930067; PubMed Central PMCID: PMCPMC3272761.

5. Battaglia M, Pesenti-Gritti P, Medland SE, Ogliari A, Tambs K, Spatola CA. A genetically informed study of the association between childhood separation anxiety, sensitivity to CO(2), panic disorder, and the effect of childhood parental loss. Arch Gen Psychiatry. 2009;66(1):64-71. Epub 2009/01/07. doi: 10.1001/archgenpsychiatry.2008.513. PubMed PMID: 19124689.

6. D’Amato FR, Zanettini C, Lampis V, Coccurello R, Pascucci T, Ventura R, et al. Unstable maternal environment, separation anxiety, and heightened CO2 sensitivity induced by gene-by-environment interplay. PLoS One. 2011;6(4):e18637. Epub 2011/04/16. doi: 10.1371/journal.pone.0018637. PubMed PMID: 21494633; PubMed Central PMCID: PMCPMC3072999.

7. Spatola CA, Scaini S, Pesenti-Gritti P, Medland SE, Moruzzi S, Ogliari A, et al. Gene-environment interactions in panic disorder and CO(2) sensitivity: Effects of events occurring early in life. Am J Med Genet B Neuropsychiatr Genet. 2011;156B(1):79-88. Epub 2010/12/25. doi: 10.1002/ajmg.b.31144. PubMed PMID: 21184587.

8. Cittaro D, Lampis V, Luchetti A, Coccurello R, Guffanti A, Felsani A, et al. Histone Modifications in a Mouse Model of Early Adversities and Panic Disorder: Role for Asic1 and Neurodevelopmental Genes. Sci Rep. 2016;6:25131. Epub 2016/04/29. doi: 10.1038/srep25131. PubMed PMID: 27121911; PubMed Central PMCID: PMCPMC4848503.

9. Battaglia M, Rossignol O, Bachand K, D’Amato FR, De Koninck Y. Amiloride modulation of carbon dioxide hypersensitivity and thermal nociceptive hypersensitivity induced by interference with early maternal environment. J Psychopharmacol. 2018:269881118784872. Epub 2018/07/04. doi: 10.1177/0269881118784872. PubMed PMID: 29968500.

10. Drug Label Information, Amiloride Hydrochloride Tablet. (https://dailymed.nlm.nih.gov/dailymed/drugInfo.cfm?setid=e0cc2d44-436a-47e8-a890-589882fff4c4) [cited 2020 01/09/2020].

11. Vidt DG. Mechanism of action, pharmacokinetics, adverse effects, and therapeutic uses of amiloride hydrochloride, a new potassium-sparing diuretic. Pharmacotherapy. 1981;1(3):179-87. Epub 1981/11/01. doi: 10.1002/j.1875-9114.1981.tb02539.x. PubMed PMID: 6927605.

12. Chapman CD, Frey WH, 2nd, Craft S, Danielyan L, Hallschmid M, Schioth HB, et al. Intranasal treatment of central nervous system dysfunction in humans. Pharm Res. 2013;30(10):2475-84. Epub 2012/11/09. doi: 10.1007/s11095-012-0915-1. PubMed PMID: 23135822; PubMed Central PMCID: PMCPMC3761088.

13. Djupesland PG, Messina JC, Mahmoud RA. The nasal approach to delivering treatment for brain diseases: an anatomic, physiologic, and delivery technology overview. Ther Deliv. 2014;5(6):709-33. Epub 2014/08/05. doi: 10.4155/tde.14.41. PubMed PMID: 25090283.

14. Corrigan M, Wilson SS, Hampton J. Safety and efficacy of intranasally administered medications in the emergency department and prehospital settings. Am J Health Syst Pharm. 2015;72(18):1544-54. Epub 2015/09/09. doi: 10.2146/ajhp140630. PubMed PMID: 26346210.

15. Ozsoy Y, Gungor S. Nasal route: an alternative approach for antiemetic drug delivery. Expert Opin Drug Deliv. 2011;8(11):1439-53. Epub 2011/10/19. doi: 10.1517/17425247.2011.607437. PubMed PMID: 22004793.

16. Bioanalytical Method Validation: Guidance for Industry. Center for Drug Evaluation and Research (CDER), Food and Drug Administration, (https://www.fda.gov/media/70858/download). accessed on March 5 2020. 2018.

17. International Conference on Harmonisation. Technical requirements for registration of pharmaceuticals for huam use. Validation of analytical procedures: text and methodology Q2(R1) (November 2005). https://database.ich.org/sites/default/files/Q2_R1__Guideline.pdf (accessed 2020 Jan 15).

18. Microbiological Tests, (71) Sterility Tests. United States Pharmacopeia 42-National Formulary 37 (USP42-NF37 2S)

19. Abdel-Hay MH, Ragab, M.A.A, Ahmed, H.H., Mohyeldin, S.M. The use of Arrhenius kinetics to evaluate different hydrolytic stability of amiloride hydrochloride and cyclopenthiazide using chromatographic methods. Microchemical Journal. 2019;147:682–90.

20. Li YN, Moore DE, Tattam BN. Photodegradation of amiloride in aqueous solution. Int J Pharm. 1999;183(2):109-16. Epub 1999/06/11. doi: 10.1016/s0378-5173(99)00035-6. PubMed PMID: 10361161.

21. FDA Guidance for industry - Nasal Spray and Inhalation Solution, Suspension, and Spray Drug Products — Chemistry, Manufacturing, and Controls Documentation. https://www.fda.gov/media/70857/download

